# Antagonistic action of a nuclear hormone receptor pair coordinates a switch from lytic to biotrophic effector production in a plant-parasitic nematode

**DOI:** 10.64898/2026.06.10.731300

**Authors:** Anika Damm, Victor Hugo Moura de Souza, Anna Dickinson, Priya Desikan, Alexis L. Sperling, George Harpum, Paul Brett, Beth Molloy, Clement Pellegrin, Chongjing Xia, Sebastian Eves-van den Akker

## Abstract

Pathogens secrete overlapping and sequential waves of effectors to manipulate their host, and yet the regulators that conduct the ensemble are poorly understood. Here, we identify the Dorsal Gland Regulator DGR-1 in the beet cyst nematode *Heterodera schachtii*. DGR-1 controls the expression of 131 putative effectors, acting as a dual-functional switch that “switches off” early stage effectors involved in plant invasion, and “switches on” later stage effectors associated with biotrophic establishment in the host. Interestingly, DGR-1 works antagonistically with the only other known transcriptional regulator of effectors in plant-parasitic nematodes, the Subventral Gland Regulator-1 (SUGR-1), to coordinate this apparent switch from lytic to biotrophic effector production. Together, DGR-1 and SUGR-1 control the expression of nearly one half of all *H. schachtii* early-stage effectors, and over one fifth of effectors of any kind. The requirement to activate lytic effector functions in the cortex, and biotrophic effector functions in the vascular cylinder, suggests that this transcription factor pair must differentially respond to signals from the host. Consistent with this, we find that diffusates from the roots of *A. thaliana* Casparian strip mutant *myb36/sgn3*, which are enriched in molecules ordinarily restricted to the vascular cylinder, upregulate *dgr-1*, but not *sugr-1*, compared to Col-0. Taken together, these data indicate that *H. schachtii* responds to compartmentalised host-derived signals to appropriately regulate spatiotemporal effector expression during infection. Given that misregulating DGR-1 results in delayed development of parasitic nematodes in Arabidopsis and Mustard, strategies which impair effector regulation as a whole may hold promise for crop protection against these agriculturally important pathogens.

## Introduction

A fundamental requirement of host-microbe interactions is the secretion of effectors by the invader to manipulate the host^1^. The action, or inaction, of effectors enables host invasion^2^, suppression of the host immune system^3^, and manipulation of cellular processes^4,5^ - and ultimately defines pathogen virulence.

Effector deployment is precisely regulated in time and space^6^. Congruent with discrete stages of infection, overlapping but distinct waves of effectors fulfil the varying requirements of the pathogen at different times^7–10^. In plant-parasitic cyst nematodes, devastating pathogens of global agricultural importance that can cause yield losses of up to 90%^11^, this is most acutely represented by a division of parasitism in two phases: a destructive “lytic” phase when the parasite enters and migrates through its host; and a “biotrophic” phase when the nematode establishes and maintains a permanent feeding site inside its host through the redifferentiation of existing cells^12^. Accordingly, effectors produced during the lytic phase are, for example, involved in the breakdown of plant cell walls^13^, while biotrophic phase effectors often suppress plant immunity or facilitate developmental reprogramming of plant tissues^14^. These different sets of effectors are predominantly produced in two different sets of specialised secretory gland cells. The subventral gland cells are principally active during the lytic phase, while the dorsal gland cell becomes active during the biotrophic phase of infection^12^.

The *subventral gland regulator 1 (sugr-1)* in the beet cyst nematode *Heterodera schachtii* is part of an expanded family of Nuclear hormone receptors (NHRs)^16^ and a master regulator of subventral gland cell effectors during the lytic phase. *sugr-1* transcription positively responds to host signals, termed effectostimulins, and in turn activates a variety of effectors involved in host invasion. These effectors facilitate breakdown of plant cell walls, thereby releasing more effectostimulins in what is proposed as a positive feedback loop driving the earliest stages of infection^15^. How this virtuous cycle is broken, and the switch from lytic to biotrophic effector production is coordinated, is unknown. Here we address this gap, with the identification of the Dorsal Gland Regular 1 DGR-1, and show that the switch in effector expression from lytic to biotrophic phases is coordinated by an antagonistic pair of nuclear hormone receptors.

## Results

### The homoplastic evolution of candidate effector regulators

To identify potential regulators of stage-specific effector expression we inferred a phylogeny of *Heterodera schachtii* nuclear NHRs (Figure 1A). Based on established precedent for *sugr-1*, we computed distance correlation coefficients between individual NHRs and effector gene expression^17^, using life-stage-specific transcriptome data^7^. The sporadic incidence of effector-connection hotspots across the phylogeny suggest widespread homoplasy of this trait.

**Figure 1:**
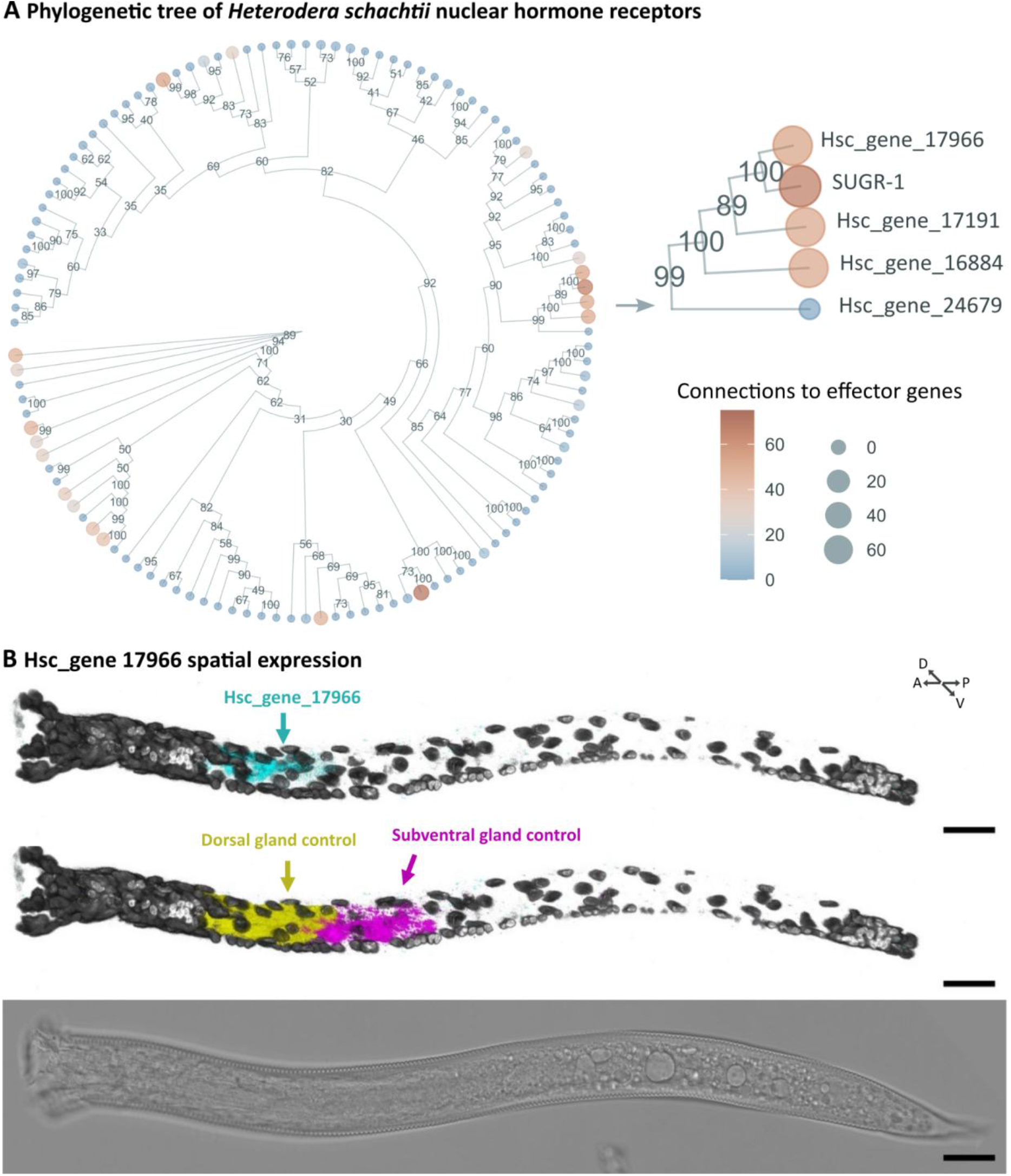
Identification of candidate effector regulators. **A)** Phylogenetic tree of all *Heterodera schachtii* nuclear hormone receptors. Colour and node size represent the number of connections to effector genes, defined as correlations in gene expression of 0.975 or above. **B)** 3D projection of a multiplexed HCR *in situ* for Hsc_gene_17966 transcripts (upper Panel, cyan), compared to dorsal gland (Hsc_gene_2729) and subventral gland (Hsc_gene_21727 *eng2*) control transcripts (middle Panel, yellow, and magenta, respectively). Nuclei stained with DAPI are shown in grey scale. Brightfield is shown in the Bottom Panel. (Scale bars, 20 μm.)

**Figure S1:**
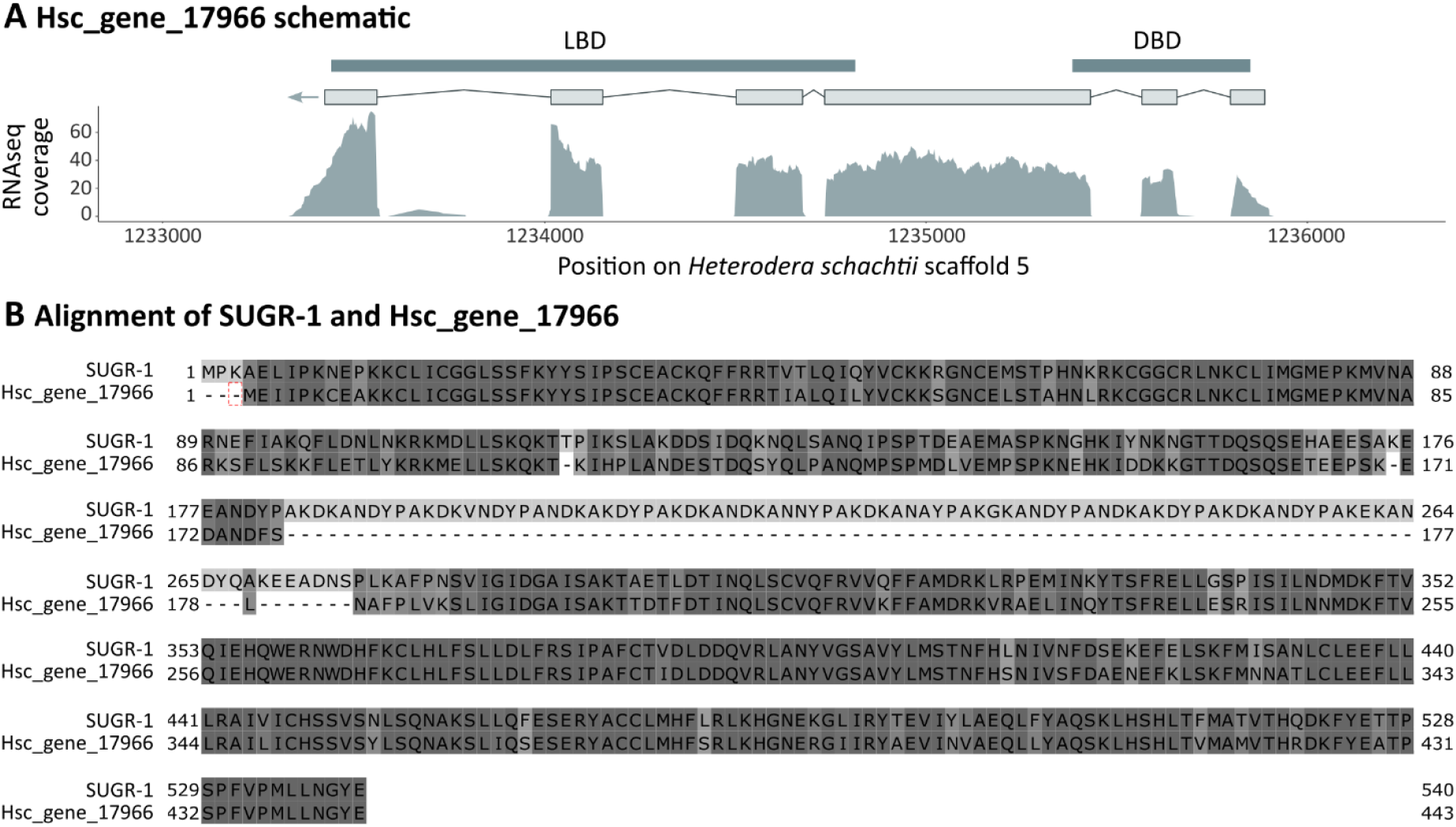
Hsc_gene_17966. **A)** Hsc_gene_17966 gene model with predicted N-terminal DNA binding domain (DBD) and C-terminal ligand binding domain (LBD). **B)** Amino acid sequence alignment of SUGR-1 (Hsc_gene_14352) and Hsc_gene_17966.

Notably, the clade containing the SUbventral Gland Regulator SUGR-1, a known regulator of early-stage effectors^15^, is exceptional in that it contains several other highly-connected NHRs. The closest homologue to SUGR-1 is Hsc_gene_17966, a canonical NHR with DNA binding and ligand binding domains (Figure S1A). SUGR-1 and Hsc_gene_17966 share 79.91 % sequence identity, with Hsc_gene_17966 lacking a ∼100 amino acid long region between DNA binding and ligand binding domain (Figure S1B). To determine if Hsc_gene_17966 and SUGR-1 could regulate similar processes, we used Sperling prep. fluorescence *in situ* hybridization chain reaction (HCR)^18^ to show that, unlike *sugr-1*, Hsc_gene_17966 is specifically expressed in the dorsal gland cell (Figure 1B).

### A pair of antagonistic transcription factors controls the switch from lytic to biotrophic effector production

To identify the function of Hsc_gene_17966, we used RNA interference-coupled RNAseq (Figure S2A+B) to show that Hsc_gene_17966 controls the expression of 1,233 genes (n = 3, |log2FC| ≥ 0.5, and *P*adj ≤ 0.001) and 131 effectors. This is a dramatic enrichment of effectors, compared to the genome as a whole (131/1,233 vs 717/26,739, p = 5.52969E-43, hypergeometric test; Figure S2C). In conjunction with the spatial expression of Hsc_gene_17966 in the dorsal gland, we have named Hsc_gene_17966 the Dorsal Gland Regulator-1 (DGR-1).

In contrast to SUGR-1, a predominantly positive regulator of cell-wall degrading, lytic phase effectors (Figure 2A), DGR-1 is a dual-functional transcription factor that both activates and represses effector gene expression (Figure 2B). DGR-1 repressed effectors are predominantly expressed during the early stages of the nematode life cycle (Figure 2B) and overlap largely with SUGR-1 activated effectors. In total 53.7 % of SUGR-1 activated genes are DGR-1 repressed (Figure S2B; hypergeometric enrichment p=6.9682E-133 (Figure S2C)). Accordingly, DGR-1 repressed effectors are enriched in GO terms associated with plant invasion like hydrolase activity or pectate lyase activity (Figure S2D).

**Figure 2:**
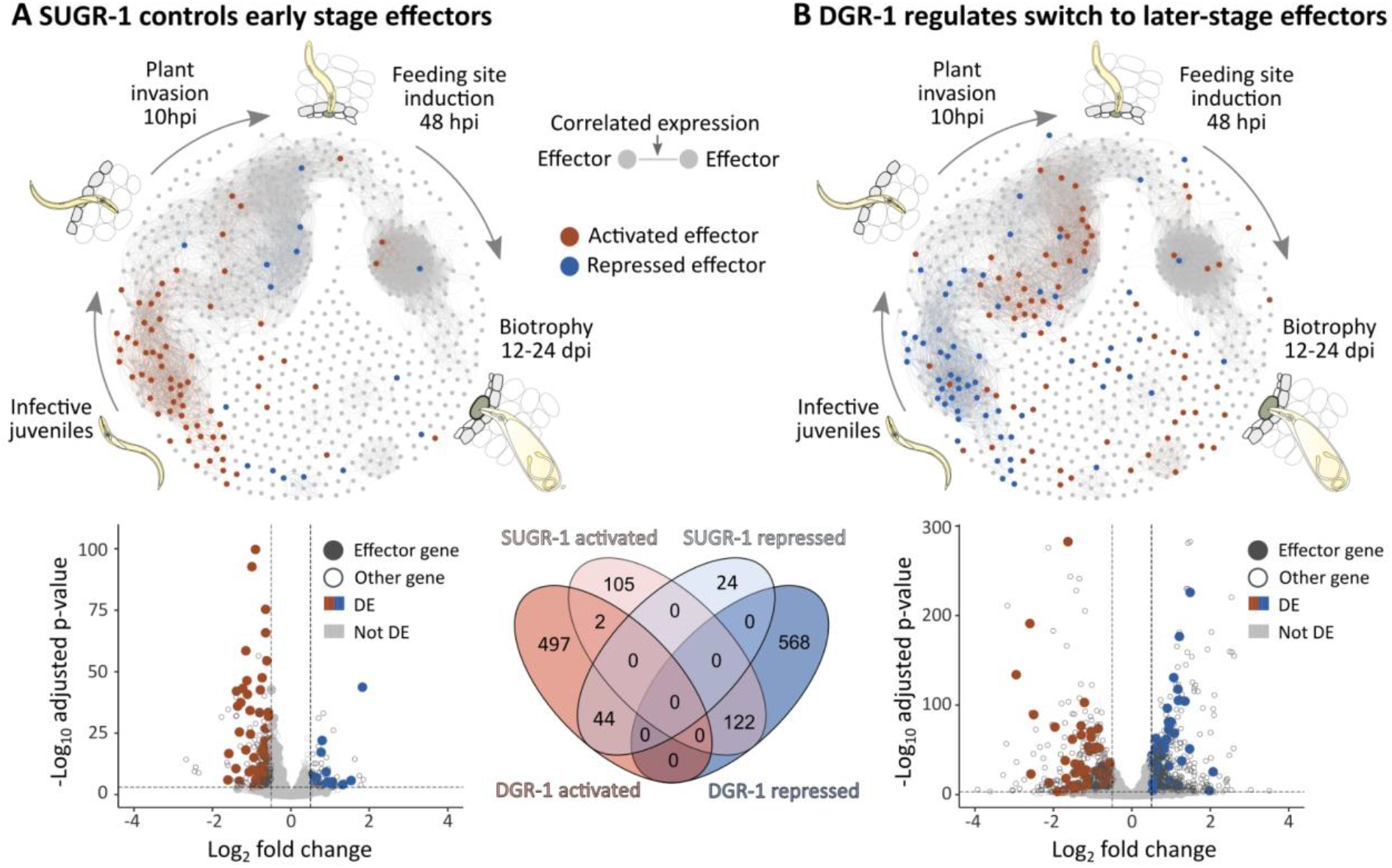
The SUGR-1/DGR-1 transcription factor duet controls a switch in effector production. Transcriptional effector networks where nodes are effector genes and edges represent correlations in gene expression across the *Heterodera schachtii* life cycle computed from publicly available life-stage transcriptomic data^7^. Volcano plots show *H. schachtii* gene expression following *sugr-1* (data from ref) **(A)** or *dgr-1* **(B)** knockdown vs. a *gfp* control. Differentially expressed (DE) effector genes (n = 3; |log2FC| ≥ 0.5, and Padj ≤ 0.001) are highlighted in red (activated) or blue (repressed). Venn diagram shows mutual antagonism for SUGR-1 and DGR-1 regulated genes.

**Figure S2:**
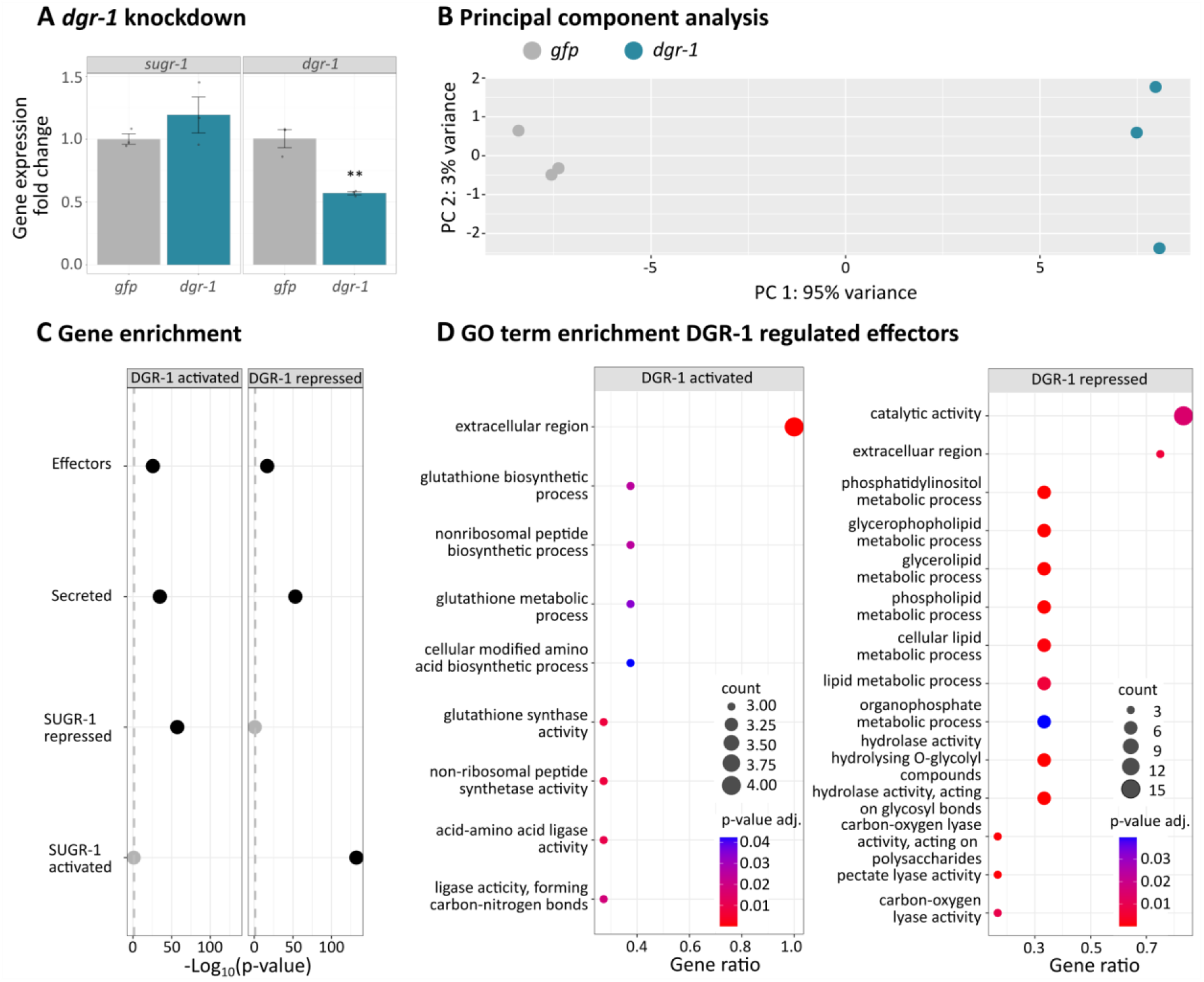
Knockdown of *dgr-1*. **A)** *sugr-1* and *dgr-1* gene expression following RNAi-mediated knockdown of *dgr-1* compared to a *gfp* control. **B)** Principal component analysis. **C)** Hypergeometric enrichment of gene sets in DGR-1 regulated genes. **D)** GO term enrichment for DGR-1 activated and repressed effector genes.

Conversely, DGR-1 activated effectors are predominantly expressed during later nematode life stages (Figure 2B) and 64.7 % of SUGR-1 repressed genes are DGR-1 activated (Figure S2B; hypergeometric enrichment p=5.68717E-58 (Figure S2C)). Consistent with promoting the biotrophic phase of infection, DGR-1 activated effectors are enriched in GO terms associated with regulation of the redox state (Figure S2D) and include known regulators of plant immunity like SPRYSECs^19^.

Taken together, these results show that the switch in effector expression from lytic to biotrophic phases is coordinated by an antagonistic pair of nuclear hormone receptors.

### A conserved DNA motif is associated with antagonistically regulated genes

DGR-1-mediated control of effector gene expression could be facilitated via direct binding to a conserved DNA motif and/or could involve downstream signalling and interaction partners. To investigate whether DGR-1 regulated genes (activated and/or repressed) are associated with a conserved DNA motif, we employed a differential motif discovery algorithm. Promoters of DGR-1 repressed genes are consistently enriched in similar motifs, broadly of the consensus sequence ATGCACTT (Figure 3A). Consistent with antagonistic action, the motif identified here as associated with DGR-1 repressed genes was previously identified as associated with SUGR-1-activated effectors^15^. To further validate the motif, we scanned all *H. schachtii* promoters for occurrences of this motif and identified 399 promoters with a perfect match. Downstream genes were significantly enriched in DGR-1 repressed and SUGR-1 activated genes as well as effectors, secreted proteins, and early-stage effectors. Later-stage effectors as well as DGR-1 activated and SUGR-1 repressed genes were not significantly enriched (Figure 3B). Furthermore, the motif was not found, indeed no clear consensus motif was found, enriched in the promoters of DGR-1 activated genes.

**Figure 3:**
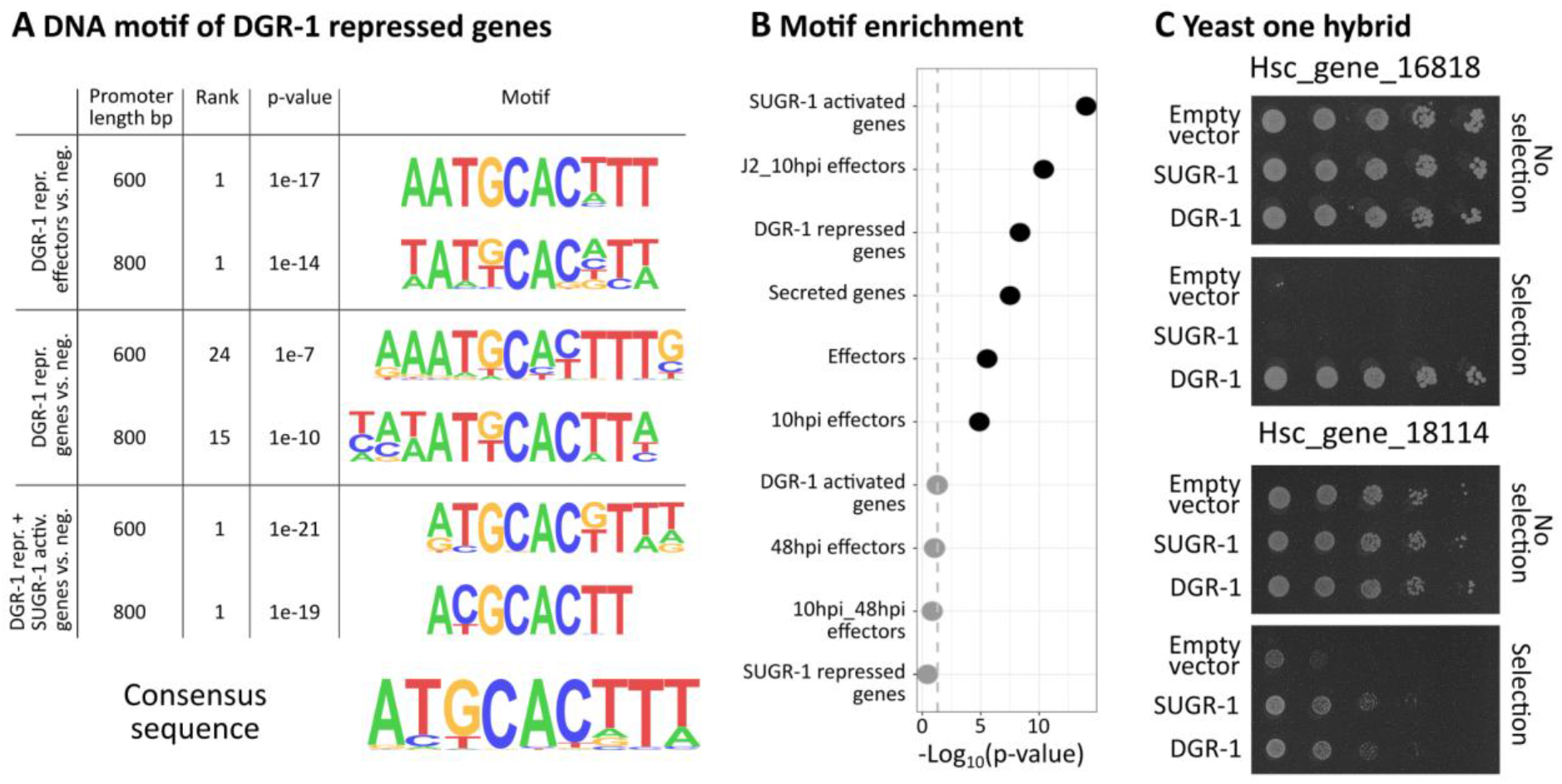
DNA motif associated with DGR-1 repressed genes. **A)** Motif enrichment analyses where respective promoters (800 bp or 600 bp from start codon) were compared to a negative control set (neg). Rank indicates order of priority in the given comparison and the p-value statistical significance of a motif’s enrichment. **B)** Hypergeometric enrichment tests for genes downstream promoters with a perfect match to ATGCACTTT. **C)** Yeast-one-hybrid screen. Yeast were grown in 1:5 serial dilutions on SD medium with (selection) or without (no selection) Aureobasidin A. Pictures are from the same screen shown in ref.^15^ but display additional information about DGR-1.

To test if DGR-1 is able to directly bind to effector promoters, we performed a yeast-one-hybrid assay with promoters of two genes that are both activated by SUGR-1 and repressed by DGR-1 and that had previously been screened for SUGR-1 binding^15^. DGR-1 was able to bind to both promoters in yeast, indicating that DGR-1 may be capable of directly mediating gene expression in *cis* for its repressive role (Figure 3C). The absence of evidence for a DGR-1 activating motif is of course not evidence of absence, but it may indicate a more complicated hierarchy of indirect action for this side of DGR-1 function.

### Knockdown of *dgr-1* affects nematode development in two hosts

Given that knockdown of *dgr-1* both upregulates lytic phase effectors and downregulates biotrophic phase effectors (Figure 2B), we investigated the effect on nematode parasitism of two host species, *Arabidopsis thaliana* and *Sinapis alba*. Interestingly, and in line with this dual-function, downregulation of *dgr-1* (Figure 4A) significantly increased the total number of nematodes per Arabidopsis plant (Figure 4B, p= 0.00011, Wilcoxon Rank Sum test), but decreased their overall fitness. *dgr-1-*silenced J2s show signs of slower development, with a significantly increased proportion of J3s compared to a *gfp* control (Figure 4B, p = 3.49E-08, Wilcoxon Rank Sum test), as well as a significantly reduced proportion of females (Figure 4B, p = 5.82E-06, Wilcoxon Rank Sum test). To evaluate the long-term fitness cost we measured female size at 14 dpi and 40 dpi and counted the number of encysting females at 92 dpi. At 40 dpi *dgr-1* silenced females are significantly smaller than control females (Figure 4C, p=0.000996, t-test). Furthermore, *dgr-1* silenced nematodes encyst significantly earlier (Figure S3A, p=0.00379, Wilcoxon Rank Sum test), potentially indicative of a stress response. Taken together, these data may indicate a reduced ability of *dgr-1-*silenced nematodes to establish a feeding site and/or properly manipulate host processes to their benefit. An infection assay in white mustard (*S. alba)* could replicate the differences in total nematode number (1/2 repeats; Figure 4D + S3B) and developmental progress (2/2 repeats; Figure 4D + S3B). While it is known that inoculation density affects nematode parasitism (Steele 1975), the second repeat in white mustard (*S. alba)* showed no difference in initial infectivity but replicated the difference in developmental progress (Figure 4D, p=0.003, Wilcoxon Rank Sum test).

**Figure 4:**
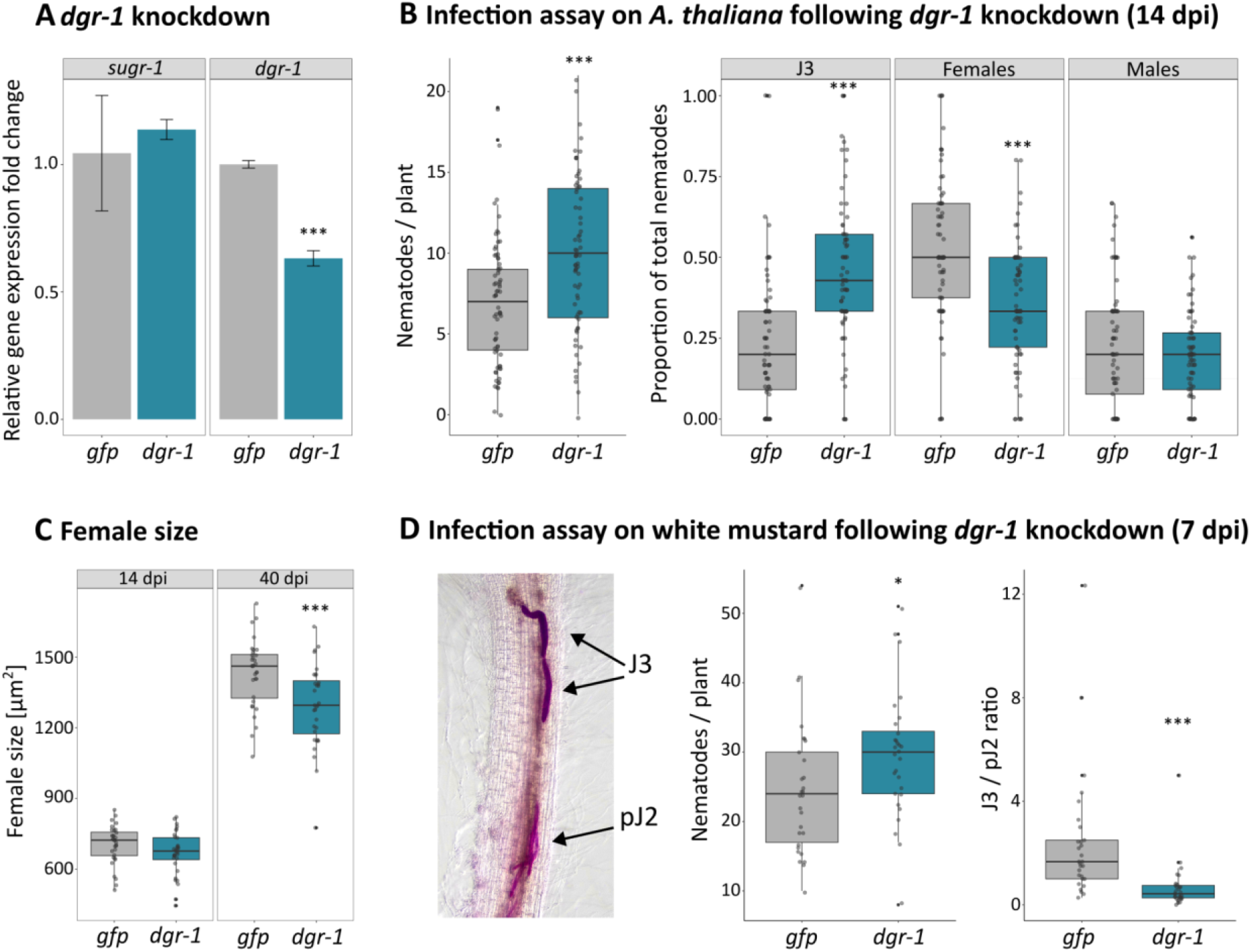
Knockdown of *dgr-1* affects nematode development and infectivity. **A)** Gene expression following knockdown of *dgr-1* compared to a *gfp* control. Data were analysed using a two-sided t-test. **B)** Infection assay on *A. thaliana* (14 dpi) following *dgr-1* knockdown. **C)** Female size at 14 dpi and 40 dpi. **D)** Infection assay on white mustard (7 dpi) following *dgr-1* knockdown. Data were analysed using a Wilcoxon Rank Sum test or two-sided t-test (Female size). Asterisks indicate significant differences to the *gfp* control (**P<0.01; ***P<0.001).

**Figure S3:**
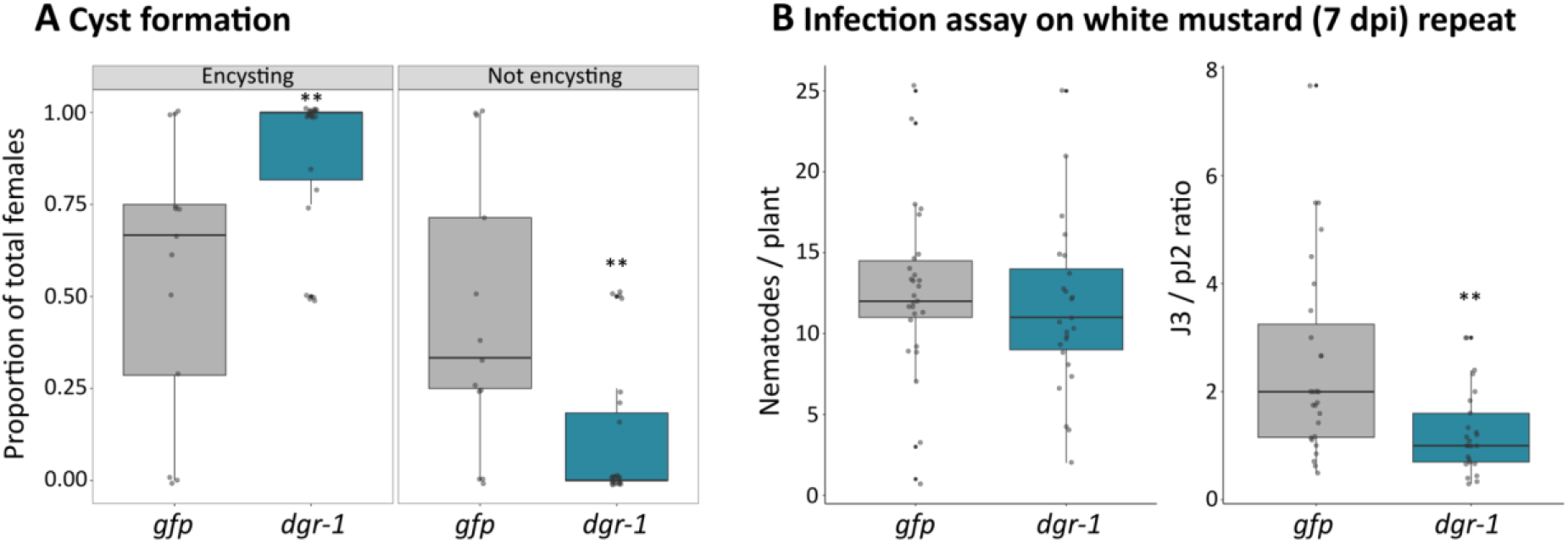
*dgr-1* knockdown phenotype. **A)** Number of encysting nematodes at 92 dpi after *dgr-1* knockdown. **B)** Repeat of infection assay on white mustard (7 dpi) following *dgr-1* knockdown. Data were analysed using a Wilcoxon Rank Sum test. Asterisks indicate significant differences to the *gfp* control (**P<0.01).

### The switch in effector deployment is linked to compartmentalisation of effectostimulins within plant tissues

These findings raise the question of where and how the *sugr-1/dgr-1* antagonistic pair is itself regulated, *in vivo*, during parasitism. Similar *to sugr-1, dgr-1* expression peaks when the nematode is inside the host at 10 hours post infection (hpi; Figure 5A). However, progressing from the juvenile (J2) stage to 10 hpi results in the ratio of *dgr-1/sugr-1* expression inverting: *dgr-1* becomes 3 times more highly expressed than *sugr-1* at 10 hpi (Figure 5B). Such a switch is likely fine-tuned by discrete plant-signals (effectostimulins). Like *sugr-1, dgr-1* expression is activated by root extract of the host plants white mustard and *Arabidopsis thaliana* but not by the non-hosts tomato and rice (Figure 5C). Fractionation of mustard extract by HPLC (Figure 5D) revealed “common” fractions, activating both transcription factors) and “unique” fractions, activating predominantly one regulator (Figure 5D).

**Figure 5:**
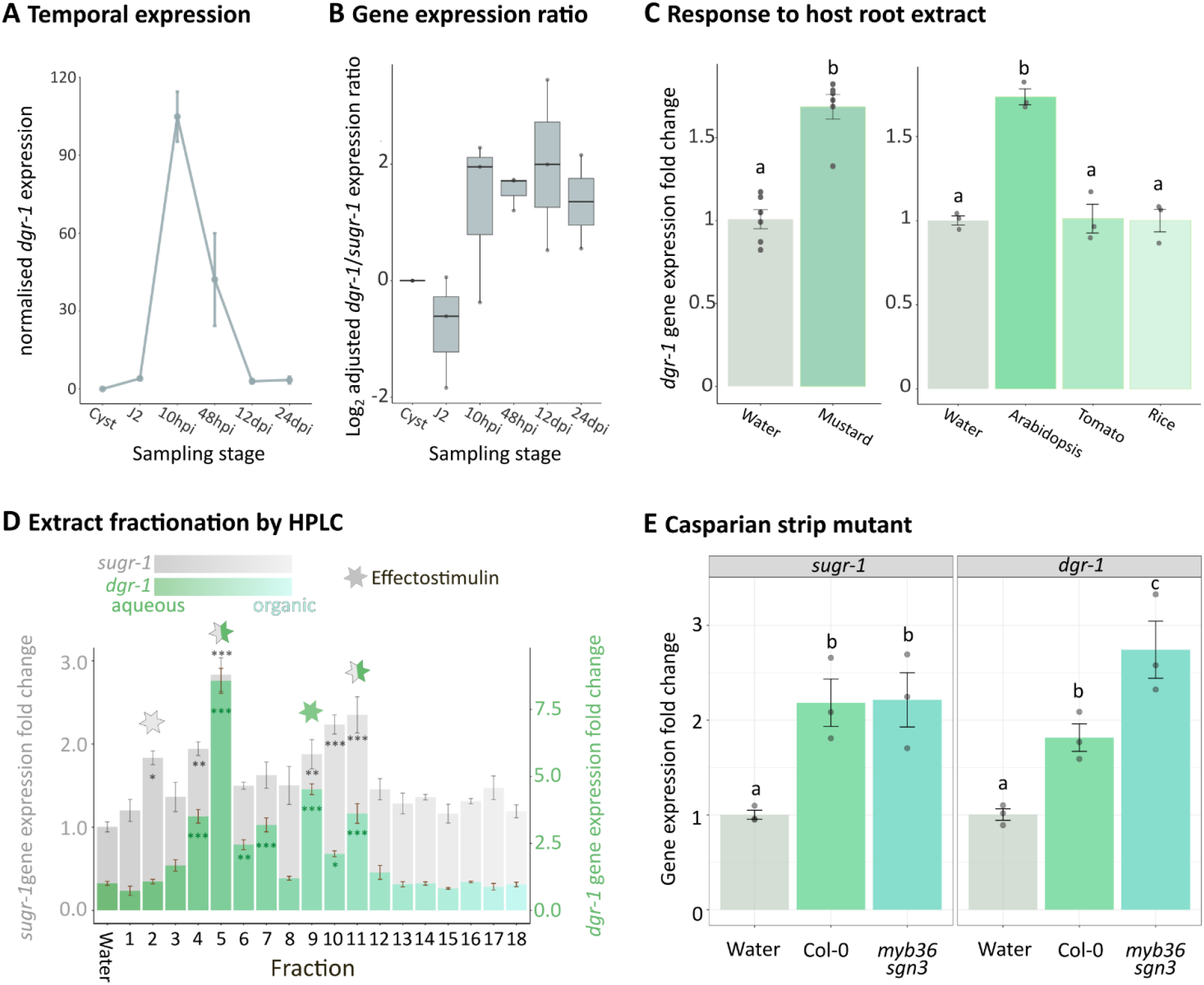
Effectostimulins are compartmentalised within the plant. **A)** *dgr-1* expression across the *Heterodera schachtii* life cycle. Data from ref.^7^ **B)** *dgr-1* / *sugr-1* gene expression ratio across the *H. schachtii* life cycle. Data from ref.^7^ **C)** Effect of plant root extract on *dgr-1* gene expression. **D)** Effect of white mustard extract HPLC fractions on *sugr-1*^15^ and *dgr-1*. Asterisks indicate significant differences to the water control (*P < 0.05; **P < 0.01; ***P < 0.001; Tukey HSD). **E)** Comparison of Casparian mutant *myb36 sgn3* diffusate with wild type Col-0 plants. In **C** and **E** treatments with the same letter are not statistically significantly different at p < 0.05 (Tukey HSD).

Given that lytic subventral gland functions need to be repressed, and biotrophic dorsal gland functions need to be promoted, in the vascular cylinder^5^, we compared the ability of wild type root diffusates to activate *dgr-1* expression with that of the Casparian strip mutant (*myb36/sgn3*^20^). The *myb36/sgn3* diffusate activated *dgr-1* significantly more than Col-0 (Figure 5E), indicating potential leakage of effectostimulins ordinarily compartmentalised in the vascular cylinder in the wild type state. No difference could be observed for *sugr-1* (Figure 5E), suggesting that nematodes hijack the plant’s native segregation of small molecules within its root tissues to tailor their parasitism gene expression appropriately.

## Discussion

Pathogens rely on the precise spatiotemporal regulation of effector gene expression^6^ and yet the corresponding regulators are poorly understood. Here, we discover Dorsal Gland Regulator DGR-1, a nuclear hormone receptor expressed in the dorsal gland cell. DGR-1 works antagonistically with the only other known transcriptional regulator of effectors in plant-parasitic nematodes, SUGR-1, to coordinate the apparent switch from lytic to subsequent biotrophic effector production (ref Molloy 2024). Thereby, DGR-1 represents not just an understanding of spatial expression in the dorsal gland, but also temporal regulation of different effector waves. Together, SUGR-1 and DGR-1 control nearly one half of all *Heterodera schachtii* effectors expressed before 48 hours post infection, and over one fifth of effectors of any kind. Thereby, SUGR-1 / DGR-1 signalling can explain, in large part, the regulation of parasitism in the beginning of the *H. schachtii* life cycle.

While SUGR-1 was previously reported as a predominantly positive regulator of gene expression (77% of DEGS up vs 23% down), in the context of DGR-1 we now appreciate that the functions of these two genes are most easily understood as a pair. So while DGR-1 is different in that it roughly equally activates and represses (44% of DEGS up vs 56% down), it is the mutually antagonistic action of both that unites them: more than half of the SUGR-1 activated genes are DGR-1 repressed and more than half of SUGR-1 repressed genes are DGR-1 activated. It is particularly striking that in the broader context of nuclear hormone receptors of this species, these two regulators of effectors are adjacent in the phylogeny and yet functionally antagonistic.

Intriguingly, DGR-1 appears to repress subventral gland lytic effectors without affecting *sugr-1* gene expression itself. The fact that the promoters of DGR-1 repressed genes are enriched in the SUG-box (ATGCACTT) is intuitive. This, in combination with the finding that DGR-1 can bind promoters of SUGR-1 activated effectors in yeast, suggests that DGR-1 might directly repress SUGR-1 activated effector genes. Interestingly, a version of the same motif (TGCACTT; the Mel-DOG box) was shown to be associated with dorsal gland effectors in the root knot nematode *Meloidogyne incognita*^21^, perhaps indicating evolutionary conserved or convergent mechanisms of effector regulation for plant-parasitic nematodes which share a common ancestor over a hundred million years ago^22^.

There is an asymmetry in the antagonism between SUGR-1 and DGR-1, proportionally more of the SUGR-1 regulon is controlled by DGR-1 than vice versa. It is intuitive to appreciate the asymmetry because, presumably, nematodes must suppress “lytic subventral gland functions” (e.g. proteases, cell wall degrading enzymes, etc.) when it is promoting “biotrophic dorsal gland functions” (e.g. suppression of immunity and establishment of the developmentally-altered feeding site etc.) to a greater extent than the inverse. However, if we knew specifically how the antagonism works it could enhance our understanding. For example, if DGR-1 represses SUGR-1-activated genes locally in the dorsal gland, the observed antagonism likely serves to maintain the distinct identity of each gland cell. Alternatively, if DGR-1 represses SUGR-1-activated genes distally in the subventral gland, the observed antagonism likely represents a switch from lytic to feeding site establishment stages of parasitism. Note, these are both important functions, not mutually exclusive, and with similar implications: the parasite must suppress lytic functions when establishing the feeding site.

These findings raise the question of how these two regulators work both antithetically but yet in concert to determine parasitism *in vivo*. Both, *dgr-1* and *sugr-1*, are activated by effectostimulins in host root extract, and are predominantly expressed 10 hours post infection. Importantly, however, the ratio of *dgr-1* to *sugr-1* expression shifts notably between second stage juveniles, collected outside the host, and nematodes sampled 10 hours post infection, suggesting that the ratio of *dgr-1* to *sugr-1* might fine tune temporal effector expression. Such fine-tuning could be achieved through compartmentalisation of *dgr-1-* or *sugr-1*-specific effectostimulins within the plant. Concordantly, diffusate of the Casparian strip mutant *myb36/sgn3* activated *dgr-1* more than wild type diffusate, while *sugr-1* activation remained unchanged. These data indicate that the Casparian strip, the major apoplastic diffusion barrier in plants, may usually restrict *dgr-1-*specific effectostimulin(s) to the vascular cylinder, the site where nematodes promote biotrophic effectors. Nematodes, therefore, appear able to hijack the plant’s native segregation of small molecules within its root tissues to tailor their parasitism gene expression appropriately.

Disrupting DGR-1 would not only repress the functions DGR-1 would normally activate, but would misregulate those that it would switch off - i.e. activate lytic functions during biotrophy. Such dysregulation could be achieved through host-induced gene silencing (HIGS). As HIGS takes place within the plant, it can avoid a potential benefit from upregulating lytic effectors before invasion while replicating the developmental effects shown on white mustard and Arabidopsis. Further routes to application could be the removal of *dgr-1*-activating effectostimulins in the host e.g. via classical breeding or gene editing, or dysregulating expression of effectostimulins within the plant. In light of cyst nematodes as one of the economically most damaging groups of plant-parasitic nematodes, causing yield losses of up to 90%^11^, targeting virulence regulators like DGR-1 might raise promising opportunities for global food security.

## Material and Methods

### Common materials

*Sinapis alba* (cv. albatross), *Arabidopsis thaliana* (Columbia-0, *myb36/sgn3*), *Solanum lycopersicum* (cv. Moneymaker), *Oryza sativa* (cv. Nipponbare*), Heterodera schachtii* populations “Bonn”, originally from Germany^7^ and “IRS,” originally from The Netherlands, the *Saccharomyces cerevisiae* Y1HGold strain (Takarabio), and competent cells of the *Escherichia coli* strain DH5α were used in this study.

### Phylogenetic tree

Nuclear hormone receptor sequences were aligned using the muscle package in R (3.46.0) and 25% of the alignment was trimmed using trimal.v1.2rev59. The phylogenetic tree was constructed using IQ-TREE 1.6.12. based on 1000 bootstraps^26–28^. Subsequently, the phylogenetic tree was read into R using ape v5.8, midpoint rooting was performed with castor v1.8.2, and a plot created using ggtree v3.12.0 with colours from scico v 1.5.0. Throughout the analyses R versions 4.2.1.and 4.4.1 were used.

### DGR-1 (Hsc_gene_17966) characterisation

The DGR-1 gene model was created using the R package genemodel v1.1.0. Protein domains were predicted using InterPro^29^. The DGR-1 and SUGR-1 amino acid sequences were aligned using Clustal Omega^30^ and visualised using Jalview^31^.

### Effectostimulin extraction and application to nematodes

*Sinapis alba* (white mustard), *A. thaliana* and *S. lycopersicum* (Tomato) plants were surface sterilised with 20 % bleach solution (Parazone) and grown on wet filter (*S. alba*) or ½ MS medium (*A. thaliana, S. lycopersicum*) at 21 °C for 7 d (S. alba), or 10d (*A. thaliana, S. lycopersicum*). *O. sativa* (Rice) seeds were briefly washed with 70 % ethanol and sterilized with 3% bleach solution for 20 min and grown on ½ MS medium at 30 °C for 7 d. Subsequently, 0.5 mg/ml (0.1 mg/ml for *A. thaliana* due to thinner roots) roots were ground in ultrapure water and centrifuged at 10,000rpm for 2min to remove bigger root particles. *S. alba* extract was further centrifuged in vivaspin columns (<3 kDa MWCO; Cytiva) at 4 °C. Finally, root extract was concentrated three times using a Concentrator plus (Eppendorf) at 45 °C. To prepare *A. thaliana* diffusate, forty seeds were germinated in 500 μL of ultrapure water at 21°C, and the water (diffusate) collected after 7 d.

*Heterodera schachtii* cysts were obtained from Stichting IRS, isolated using sieves (4,000, 2,000, 500, 125, 63 microns), transferred to hatching jars (Jane Maddern Cosmetic Containers), and hatching was induced using 3 mM Zinc chloride solution. Hatching jars were kept at 21 °C, and hatched juveniles were collected every 2-3 d. At least 17,000 second stage juveniles (J2s) were treated with 50 μl extract for 4 h at 700 rpm, and subsequently flash frozen in liquid nitrogen and stored at -80 °C.

### High Performance Liquid Chromatography (HPLC)

High Performance Liquid Chromatography was performed with a Shimadzu HPLC (Shimadzu Europa GmbH) comprising Nexera X2 binary pump and autosampler with 500 μL sample loop, a Prominence column oven and diode array detector, and fraction collector, and controlled using Shimadzu’s Lab Solutions software (version 5.72). Separation was carried out with a YMC-Pack Pro C18 column, 250 mm × 10.0 mm ID S-5 μm 12 nm (YMC Europe GmbH Dinslaken). The column was maintained at 40 °C and a gradient used for elution at 4.0 mL/min flow rate. The initial composition of 95% mobile phase A (0.1% formic acid) and 5% mobile phase B (acetonitrile) was changed to 100% B over 16 min, and held isocratic at 100% B for a further four min. This was followed by returning to the initial composition over 1 min and re-equilibrating the column for a further 9 min. In total 200 μL of the effectostimulin-containing root extract was injected and fractions were collected every minute. Subsequently, fractions from eight sample runs were pooled, evaporated and resuspended in 250 μL ultrapure water.

### RNA extraction and qPCR

Frozen nematodes were ground to a fine powder in a Geno/Grinder 2010 (Spex Sample Prep) in two 30 s long cycles at 1,200 strokes/min. Subsequently, RNA was extracted using the RNeasy Plant Mini Kit (Qiagen) following the manufacturer’s instructions and using QIAshredder columns and on-column DNAse digestion. cDNA was synthesized with 400 ng RNA, oligodT15 primers (Promega) and Superscript iv (ThermoFisher) following the manufacturer’s instructions, using the optional RNAse H digestion. qPCR was performed with two reference genes as described in ref.^15^ and using the LUNA Universal qPCR Master Mix (NEB) following the manufacturer’s instructions and 1 μL of cDNA. Primer sequences for dgr-1 are GCAAGGAAAAGCTTTTTATCC (forward) and AGGCTCTTTACTAATGGAAAAGCG (reverse). Data were normalised using the Pfaffl method. The assumptions of normality and variance homogeneity were checked by visual inspection of QQ plots with standardized residuals and residuals versus fitted plots, and data analysed using one-way ANOVA and Tukey HSD multiple pairwise comparisons.

### Life-stage specific gene expression

Life-stage specific gene expression data were obtained from ref.^7^ and plotted using ggplot2. To calculate *dgr-1* / *sugr-1* gene expression ratios, count data were TPM normalised and ratios Log_2_ transformed.

### Fluorescence in situ hybridisation chain reaction

Fluorescence *in situ* hybridisation chain reaction was performed as described in ref.^18^. The probes to Hsc_gene_17966, Hsc_gene_2729, Hsc_gene_21727 (*eng2*) and *in situ* reagents were designed and purchased from Molecular Instruments, Inc. Images were taken on a Leica Stellaris 8 FALCON confocal microscope, with minor adjustments made to the brightness and contrast, and prepared in ImageJ^32^. 3D projections were created with the Leica Cyclone 3DR software. No further image manipulation was performed.

### RNA interference

Double stranded RNA against *dgr-1* and *gfp* (21 bp) was synthesised using the Silencer siRNA Construction Kit (cat. no. AM1620). Approximately 17,000 H. schachtii J2s were soaked in silencing mix (30 μl RNA, 2.5 μl Octopamine [1M], and 17 μl ultrapure water) for 48 h at 700 rpm. Subsequently, nematodes were flash frozen in liquid nitrogen and RNA extracted as described above. Knockdown of *dgr-1* compared to a *gfp* control was confirmed via qPCR. Subsequently, samples were sent for sequencing.

### RNA sequencing

RNA sequencing and library construction were performed by Novogene. mRNA libraries were made by poly-A enrichment (poly-T oligo-attached magnetic beads), fragmentation, cDNA synthesis (using random hexamer primers), end-repair, A-tailing, adapter ligation, size selection, amplification, and purification. Illumina sequencing was performed using 150 bp paired-end reads, generating 5G raw data per sample. Quality of RNA sequencing reads was checked with FastQC^33^ and trimming was performed using BBduk^34^. Subsequently, reads were mapped to the *H. schachtii* reference genome^7^ using STAR^35^, and counted using HTseq^36^. Differentially expressed genes were identified using DESeq2^37^, and pairwise comparisons of all samples (|log2FC| ≥ 0.5 and *P*adj ≤ 0.001). Volcano plots were made using EnhancedVolcano^38^, and GO term enrichment analyses were performed using the gprofiler2^39^ and enrichplot^40^.

### Effector network analyses

The manually curated effector definitions and transcriptional network of predicted *H. schachtii* effectors with an arbitrary edge threshold set at a distance correlation coefficient above 0.975 from ref.^17^ were used in this paper. DGR-1 and SUGR-1 regulated effectors were highlighted and the network visualized using Gephi v0.10.1^41^. Scripts for transcriptional network analyses can be found at: https://github.com/BethMolloy/Effectorome_H_schachtii/tree/main.

### DNA motif identification

Proximal 5′ promoter regions were defined as 600 or 800 bases of intergenic space, where available, upstream of the coding start site of all *H. schachtii* genes and were predicted using a series of custom Python scripts (https://github.com/sebastianevda/H.schachtii_promoter_regions). From this database of promoter regions, subsets were extracted and compared. Comparisons included DGR-1 activated effectors vs a random set of 666 genes; DGR-1 activated genes vs a random set of 666 genes; DGR-1 repressed effectors vs a random set of 666 genes; DGR-1 repressed genes vs a random set of 666 genes; DGR-1 repressed genes + SUGR-1 activated effectors vs a random set of 666 genes. Enriched motifs were identified using HOMER^42^. Promoters containing a perfect match to ATGCACTTT were identified using FIMO^43^ and hypergeometric enrichment tests performed.

### Yeast-one-hybrid

Yeast-one-hybrid screens (including constructs and yeast strains) were performed as described in ref.^15^. In short: yeast overnight cultures were diluted to OD600 = 0.6, plated in 1:5 serial dilution on SD medium with varying concentrations of the antibiotic Aureobasidin A. Pictures were taken after 2, 3, and 4 days. Pictures shown are from the Yeast-one-hybrid screen in^15^ but include previously hidden information about DGR-1.

### Nematode infection assays

Sterile *H. schachtii* cysts were picked from *Sinapis alba* plates and hatched as described previously. Subsequently, hatched juveniles were silenced in *dgr-1* or *gfp* as described previously.

For infection assays on *A. thaliana*, Col-0 seeds were sown on standard KNOP medium (0.8% daishin agar) in 5 cm, deep-well petri dishes. After 4 d at 4 °C, plants were grown for 21 d at 21 °C under 16h day and 8h night conditions. On day 21, plants were inoculated with 80 nematodes per plant. Fourteen days post inoculation J3s, males, and female nematodes were counted under a stereo microscope. Three repeats with 23 replicates per treatment (total 69 replicates per treatment) were performed. Furthermore, the size of 32 females per treatment was measured in ImageJ at 14 dpi and 40 dpi and the number of encysting females was counted at 92 dpi.

For infection assays on *S. alba*, white mustard seedlings were grown on 1.5% daishin agar at 21 °C for 5 d. On day five, plants were inoculated with 100 nematodes per plant. Acid fuchsin staining was performed 7 days post inoculation by bleaching roots for 10 min in 1.5% bleach, resuspended in water for at least 15 min, followed by boiling in acid fuchsin for 1 min and acidified glycerol for 30 s. Subsequently, roots were kept in acidified glycerol (1 mL of HCl for 1 L of glycerol), and nematodes were counted under a stereo microscope. The experiment was performed with ∼25 replicates per treatment.

Data were analysed using a Wilcoxon Rank Sum test. Female sizes were analysed using a two-sided t-test and the assumptions of normality and variance homogeneity checked using a Shapiro-Wilk test and a Levene’s test.

## Notes

### Competing Interest Statement

The authors have declared no competing interest.

